# Fe_3_O_4_ nanoparticles to optimize the co-digestion of vinasse, filter cake, and deacetylation liquor: operational aspects and microbiological routes

**DOI:** 10.1101/2022.03.21.484299

**Authors:** Maria Paula. C. Volpi, Gustavo Mockaitis, Bruna S. Moraes

**Affiliations:** Interdisciplinary Center of Energy Planning, University of Campinas (NIPE/UNICAMP), R. Cora Coralina, 330 - Cidade Universitária, Campinas - SP, 13083-896, Brazil; Interdisciplinary Research Group on Biotechnology Applied to the Agriculture and the Environment (GBMA), School of Agricultural Engineering (FEAGRI), University of Campinas (UNICAMP), Av. Candido Rondon, 501 - Cidade Universitária, Campinas - SP, 13083-875, Brazil

**Keywords:** Nanoparticles, Co-digestion, Methane optimization, 1G2G ethanol residues

## Abstract

The co-digestion of residues from the sugarcane industry has already proven to be a highly attractive process for biogas production through anaerobic digestion (AD). The use of residues such as vinasse (1G) filter cake (1G) and deacetylation liquor (2G) in CSTR operation showed the potential for integrating 1G and 2G ethanol biorefineries through AD in previous work by our research group. The use of nanoparticles (NP) is a favorable way to optimize AD processes, as these additives allow the introduction of nutrients to the process more assertively concerning the distribution and interaction with microorganisms. The present work proposed the optimization of the co-digestion of vinasse, filter cake, and deacetylation liquor in a continuous reactor by adding Fe_3_O_4_ NP, comparing the results with a previous reactor operation without NP. Initially, tests were carried out in batches with different NP concentrations, resulting in 5 mg L^-1^ as the best concentration to be added in the continuous reactor along the increments of the applied organic rate load (ORL). CH_4_ production reached a maximum value of 2.8 ± 0.1 NLCH_4_ gVS^-1^ and the organic matter removal reached 71 ± 0.9%, in phase VI (ORL of 5.5 gVS L^-1^ day^-1^). This production was 90% higher than the reactor co-digestion operation without NP. The pH and alkalinity results indicated the methanogenesis stabilization within 60 days of operation: 30 days before when there was no NP added. The AD development was stable, presenting low variations in the oxidation-reduction potential (ORP) and stable organic acid (OA) concentrations, which indicated the propionic acid route to produce CH_4_ was predominant. The main methanogenic *Archeae* identified was *Methanoculleus*, indicating that the predominant metabolic route was that of acetate (SAO) coupled with hydrogenotrophic methanogenesis. The use of Fe_3_O_4_ NP managed to improve the AD from the 1G2G ethanol production residues and stimulated the microbial community growth, not modifying the preferable metabolic pathways.

## 1. INTRODUCTION

Anaerobic digestion (AD) is a process of managing liquid and solid waste that allows energy recovery through methane (CH_4_) ^1^. This technique is used for different types of residues, and the literature has shown the great potential for CH_4_ generation from sugarcane residues, especially vinasse ^2–4^.

Within this context, anaerobic co-digestion has emerged to improve biogas production. Co-digestion is characterized by the AD of two or more substrates, which is an option to overcome the disadvantages of mono-digestion, mainly concerning the nutrient balance and improve the economic viability of AD plants ^5^. Co-digestion has the advantage of optimizing CH_4_ production, in addition to better stabilizing the process. With the presence of different substrates, synergistic effects can be provided within the reactor, increasing the load of biodegradable compounds ^5^.

Promoting the co-digestion of residues from the sugarcane industry can be an alternative to improving the management of various residues obtained in this biorefinery, in addition to increasing CH_4_ generation. Apart from vinasse, filter cake is a lignocellulosic residue obtained from ethanol production that has a high potential for biogas production ^6, 7^ and can potentiate the vinasse CH_4_ production by co-digesting these two residues ^8^. However, there is not a vast body of literature on the use of residues from 2G ethanol production for AD, mainly emphasizing only 2G vinasse exploitation ^9^ but not reporting the use of liquors generated within the 2G ethanol production process. In a study carried out by Brenelli et al. ^10^, an alkaline pretreatment of sugarcane straw was performed for 2G ethanol production. Within this process, straw deacetylation was carried out before the hydrothermal pretreatment as the straw hemicellulose is highly acetylated. The residue generated from this process, called deacetylation liquor, is rich in volatile fatty acids such as acetic acid, and formic acid, and is promising for CH_4_ production through AD or co-digestion ^7^

In previous studies carried out by our research group, the co-digestion of the sugarcane industry residues for biogas production was proposed to improve the 1G2G ethanol biorefinery integration. The results showed that the co-digestion of vinasse, filter cake, and deacetylation liquor in a Semi-continuous Stirred Tank Reactor (s-CSTR) reached 230 NmLCH_4_ gSV^-1^ and organic matter removal efficiency of 83% ± 13%, confirming that the co-digestion was able to increase CH_4_ production when compared to the vinasse mono-digestion ^8^

The literature reports that using additives can potentially improve the reactors’ operation performance concerning the CH_4_ yield, mainly related to micronutrient addition ^11^. Scherer et al. ^12^ classified the importance of micronutrients for methanogenic organisms as follows: Fe >> Zn> Ni> Cu = Co = Mo> Mn, indicating that such elements play essential roles in the construction of methanogenic cells. In addition, many of these micronutrients have concentrations that must be met, as cell growth may be limited or inhibited. Zhang et al. ^13^ showed that there is a limitation to the methanogenic microorganism growth in terms of cell density if the concentrations of Co, Ni, Fe, Zn, Cu are less than 4.8, 1.32, 1.13, 0.12 g L^-1^, respectively.

Among the different trace elements, Fe is important to stimulate citrocome and ferredoxin formation and the cellular energy metabolism, mainly of methanogenic archaea ^14^. Fe is also important to catalyze chemical reactions of some metalloenzymes used in acetogenesis, such as dehydrogenase format and carbon monoxide dehydrogenase ^14^. The hydrolysis and acidification phase of AD is also benefited by Fe as a growth factor as Fe supplementation can accelerate these steps ^15^.

In the study by Demirel and Scherer ^16^, the addition of Fe_3_O_4_ improved the biogas production and its CH_4_ content when applying cow dung and chicken litter as substrates. Zhang et al. ^17^ showed that Zerovalent Iron (ZVI) contributes to optimizing the anaerobic environment for wastewater treatment and that promotes the growth of methanogens with higher removal of chemical oxygen demand (COD).

One of the ways to promote this addition of components to optimize AD is by using nanoparticles (NPs). Nanotechnology allows the manipulation of matter on a nanoscale (1 to 100 nm), and NPs are materials found in this size range ^18^. The nano size is important because it allows greater mobility of the active compound in the environment, in addition to allowing interaction with the biological system, facilitating the passage of the compound in cell membranes, absorption, and distribution in the metabolism. This happens due to its mesoscopic effect, small object effect, quantum size effect, and surface effect and to have the greater surface area and dispersibility ^18–20^.

Some authors have already studied the use of different nanoparticles to optimize biogas production in different types of waste. Henssein et al. ^21^ studied the use of NPs in AD of poultry litter. They observed that the CH_4_ production increased with the addition of NPs, namely the NP concentrations (in mg L^-1^) of 12 Ni (38.4% increase), 5.4 Co (29.7% increase), 100 Fe (29.1% increase), and 15 Fe_3_O_4_ (27.5% increase). Mu et al. ^22^ studied the effect of metal oxide nanoparticles (nano-TiO_2_, nano-Al_2_O_3_, nano-SiO_2_, and nano-ZnO) on AD using activated sludge as substrate, and the results showed that only Nano-ZnO had an inhibitory effect on CH_4_ production in concentrations starting at 30 mg g^-1^ total suspended solids (TSS). Abdeslam et al. ^18^ used the metallic NPs Co, Ni, Fe, and Fe_3_O_4_ to compare the biogas and CH_4_ production from the AD of cattle manure and the results showed the methane yield increased significantly (p < 0.05): 2, 2.17, 1.67 and 2.16 times compared to the control, respectively. Wang et al. ^23^ investigated the effects of representative NPs, (nZVI, Fe_2_O_3_ NPs) on CH_4_ production during the AD of waste activated sludge: the concentration of 10 mg g^-1^ TSS nZVI and 100 mg g^-1^ TSS Fe_2_O_3_ NPs increased methane production to 120% and 117% in relation to the control, respectively. The literature has shown that experiments with Fe_3_O_4_, which are magnetic NPs, improved the AD process due to their characteristics of superparamagnetic, high coercivity, and low Curie temperature. In addition to these characteristics, Fe_3_O_4_ NPs are also non-toxic and biocompatible ^20, 24^, which may favor AD processes.

To date, studies on the use of NPs to optimize biogas production in co-digestion with residues from the sugarcane industry have not been found in the literature. In our previous work ^8^, the co-digestion of residues from the sugarcane industry and the characterization of the microbial community were carried out. The objective of the present study was to co-digest vinasse, filter cake, and deacetylation liquor in an s-CSTR reactor, adding Fe_3_O_4_ nanoparticles, in order to optimize the biological process. Firstly, Biochemical Methane Potential (BMP) assays were performed with different concentrations of Fe_3_O_4_ NPs to figure out the best concentration to be applied in the reactor operation. Secondly, the semi-CSTR operation was performed, as well as its microbial community characterization and a comparison was made to the inoculum to evaluate the effect on the biological process changing by adding NPs.

## 2. MATERIAL AND METHODS

### 2.1 Residues and Inoculum

The substrates included vinasse and filter cake from the Iracema sugarcane mill (São Martinho group, Iracemápolis, São Paulo state, Brazil) and the liquor from the straw pretreatment process, performed at the National Biorenovables Laboratory (LNBR) from the Brazilian Center for Research in Energy and Materials (CNPEM). Deacetylation pre-treatment was applied to sugarcane straw on a bench-scale as described in Brenelli et al. ^10^. The anaerobic consortium of the mesophilic reactor (BIOPAC®ICX - Paques) from the aforementioned Iracema mill was used as inoculum. The substrates were characterized in terms of solids series, volatile solids (VS) and total solids (TS) through method 2540, and pH (pHmeter PG 1800), according to Standard Methods - APHA ^25^, Organic acids (OA), alcohol, carbohydrates, in High-Performance Liquid Chromatography (HPLC, Shimadzu®). The HPLC consisted of a pump equipped apparatus (LC-10ADVP), automatic sampler (SIL-20A HT), CTO-20A column at 43 °C, (SDP-M10 AVP), and Aminex HPX-87H column (300 mm, 7.8 mm, BioRad). The mobile phase was H_2_SO_4_ (0.01 N) at 0.5 ml min^-1^. The inoculum was characterized in terms of VS and TS. The inoculum presented 0.0076 ± 0.00 g mL^-1^ in terms of VS and 0.0146 ± 0.00 in terms of TS. The vinasse presented 0.014 ± 0.00 g mL^-1^ of VS and 0.0176 ± 0.00 g mL^-1^ of TS, the deacetylation liquor 0.0123 ± 0.00 g mL^-1^ of VS and 0.0219 ± 0.00 g mL^-1^ of TS, and filter cake 0.5454 ± 0.53 g mL^-1^ of VS and 0.6197 ± 0.54 g mL^-1^ of TS. The pH of the inoculum was 8.57 ± 0.14, the pH of vinasse was 4.25 ± 0.17 and the deacetylation liquor the pH was 9.86 ± 0.15. The elemental composition was performed for the characterization of filter cake in the Elementary Carbon, Nitrogen, Hydrogen and Sulfur Analyzer equipment (Brand: Elementar; Model: Vario MACRO Cube - Hanau, Germany), was obtained 1.88% of N, 31.07% of C, 6.56% of H and 0.3% of S, all in terms of TS.

The characterization, in terms of OA, alcohol, and carbohydrates for liquid residues, is presented in Table 1.

**Table 1.** Characterization of OA, carbohydrates, and alcohols of liquid residues

| Compounds | Vinasse (mg L<sup>-1</sup>) | Deacetylation Liquor (mg L<sup>-1</sup>) |
| --- | --- | --- |
| Acetate | 1268.41 | 3250.00 |
| Formate | -- | 650.00 |
| Lactate | 3706.94 | 423.18 |
| Propionate | 634.85 | 368.29 |
| Butyrate | -- | 250.02 |
| Isovalerate | 931.63 | 269.03 |
| Glucose | 809.05 | 546.23 |
| Methanol | 8674.83 | -- |

--: not carried out

### 2.2 Batch Tests

Batch tests were performed for the co-digestion of residues (vinasse + filter cake + deacetylation liquor) in the proportion of 70:20:10 (in terms of VS), respectively, following our previous work ^8^ with different concentrations of Fe_3_O_4_ NPs to identify the best concentration to be used in the s-CSTR reactor. The tests were conducted in 250 ml Duran flasks, under 55 °C, in which the inoculum was acclimated initially. On the first day, the temperature was increased to 40 °C, then to 45 °C and within 4 days it reached 55 °C. The inoculum was then kept for 1 week at 55 °C, prior to the experiment start-up. The experiments were conducted in triplicate, with a 2:1 inoculum to substrate ratio (in terms of VS) added to each flask, following the protocol of Triolo et al. ^26^ and the VDI 4630 methodology ^27^. The pH of solution flasks was corrected to neutrality by adding solutions of NaOH (0.5 M) or H_2_SO_4_ (1 M) when necessary. N_2_ flowed into the headspace of each vial. The biogas produced was collected from the headspace with the Gastight Hamilton Super Syringe (1L) syringe through the flasks’ rubber septum. Gas chromatography analyzes were also carried out to detect the concentration of CH_4_ produced in the gas chromatograph (Construmaq MOD. U-13 São Carlos). The carrier gas was hydrogen (H2) gas (30 cm s^-1^) and the injection volume was 3 mL. The GC Column was made of 3-meter-long stainless steel, 1/8 inch diameter, and packaged with Molecular Tamper 5A for separation of O_2_ and N_2_ and CH_4_ in the thermal conductivity detector (TCD). Digestion was terminated when the daily biogas production per batch was less than 1% of the accumulated gas production. After the assay, the values were corrected for standard temperature and pressure (STP) conditions (273 K, 1.013 hPa).

The different concentrations of Fe_3_O_4_ NP used in each bottle are described in Table 2. The choice of concentrations was made based on studies with NP and AD that the literature shows ^18, 23^. It is worth mentioning that a control flask was made (Flaks 1-Table2), adding only the inoculum and co-digestion, without NPs, to compare with the other bottles that contained NPs, and to evaluate the optimization of the process. Analysis of variance (ANOVA) was used to identify the existence of significant differences between the treatments (p < 0.05).

**Table 2.** Design of experiments of BMP

| Flasks | Name in Graph | Fe<sub>3</sub>O<sub>4</sub> NP<br>Concentration (mg L<sup>-1</sup>) |
| --- | --- | --- |
| 1- Inoculum + Co-digestion | Control | 0 |
| 2- Inoculum + Co-digestion + NP | NP 1 | 1 |
| 3- Inoculum + Co-digestion + NP | NP 2 | 5 |
| 4- Inoculum + Co-digestion + NP | NP 3 | 10 |
| 5- Inoculum + Co-digestion + NP | NP 4 | 20 |

### 2.3 Operation of Semi-Continuous Stirred Tank Reactor (s-CSTR)

The s-CSTR operation was followed according to previous work by our research group ^8^. The 5L-Duran flask with 4L-working volume kept under agitation at 150 rpm by using an orbital shaking table Marconi MA 140. The operating temperature was 55°C, maintained by recirculating hot water through a serpentine. The reactor was fed once a day with a blend of co-substrates (in terms of volatile solids, VS): 70% of vinasse, 20% of filter cake, and 10% of deacetylation liquor, totaling 33.45 gVS L^-1^. The Organic Load Rate (OLR) was increased throughout the operation to reach the maximum OLR without the reactor collapsing. Fe_3_O_4_ NP was added prior to each OLR increase, which occurred only after reactor stabilization in the respective OLR, in terms of CH_4_ production. Table 3 presents the operational parameter values applied to the s-CSTR according to the respective operation phases and the days on which Fe_3_O_4_ NPs were added.

**Table 3.** Reactor operation phases and the respective applied ORLs, feeding rate flows, and HRT.

| Phase in Graph | OLR (gVS L<sup>-1</sup> day<sup>-1</sup>) | Feeding rate (L day<sup>-1</sup>) | HRT (days) | NP Addition Day |
| --- | --- | --- | --- | --- |
| I | 2 | 0.250 | 16 | 24 |
| II | 2.35 | 0.285 | 14 | 47 |
| III | 3 | 0.363 | 11 | 72 |
| IV | 4 | 0.500 | 8 | 95 |
| V | 4.70 | 0.571 | 7 | 109 |
| VI | 5.5 | 0.666 | 6 | 123 |
| VII | 6.6 | 0.800 | 5 | 136 |
| VIII | 8 | 1.000 | 4 | 150 |
| IX | 9 | 1.140 | 3.5 | -- |

Note: --: not added

#### 2.3.1 s-CSTR monitoring analyses

The volume of biogas produced was measured using the Ritter gas meter, Germany. The CH_4_ content was determined by gas chromatography (Construmaq-MOD U-13, São Carlos) five times a week. OA, carbohydrates, alcohols, and organic matter content (in terms of VS) in the digestate were monitored following the same methodology described in the characterization of residues (Section 2.1). The alkalinity from digestate also was determined using the titration method APHA, ^25^. The pH and the Oxidation-reduction potential (ORP) of digestate were measured, immediately after sampling (before feeding) using a specific electrode for Digimed ORP. The pH was monitored also in the feed. All reactor monitoring analyses followed Volpi et al. ^8^.

### 2.4 Molecular Analysis in Biological Studies

The microbial community of the inoculum was identified before being inserted in the reactor Sample A1, and after CH_4_ production was stabilized in the OLR of 4 gVS L^-1^ day^-1^ (Sample A2), to evaluate the change in the microbial community with the changes of the metabolic routes for the CH_4_ production and with the addition of Fe_3_O_4_ NP. The extraction and quantification and sequencing protocol were followed as described in Volpi et al. ^8^. Raw sequences were deposited in BioSample NCBI under accession number BioProject ID PRJNA781620.

### 2.5 NP preparations and characterization

Fe_3_O_4_ NP was used due to the better performance of these NP in AD according to the literature ^18, 28, 29^. The Fe_3_O_4_ NP used were IRON (II, III) OXIDE, NANOPOWDER, 50-100 N-SIGMA-ALDRICH. They were then diluted in distilled water at pH 7, in a glass bottle. Sodium dodecylbenzene sulfonate (SDS) at 0.1mM was used as a dispersing reagent to ensure NPs dispersion before use, as SDS has been shown to not significantly affect CH_4_ production ^21, 23^. To characterize the size of these NPs, an analysis was performed on the Laser Diffraction Particle Size Analyzer - MASTERSIZER-3000 (MALVERN INSTRUMENTS-MAZ3000-Worcestershire, U.K.). Measurements were taken in Wet Mode - HIDRO EV. The mathematical model used: Mie. It considers that the particles are spherical and that they are not opaque - thus taking into account the diffraction and diffusion of light in the particle and the medium. They were made for samples of pure NP.

## 3. RESULTS AND DISCUSSION

### 3.1 Characterization of Fe_3_O_4_ NP

Figure 1 shows the size and distribution of Fe_3_O_4_ NP diluted in water pH 7. Figure 1a shows two populations, one up to nano size (0.1 µm) and the other that starts from 0.3 µm and is not considered a nanoparticle.

**Figure 1.**
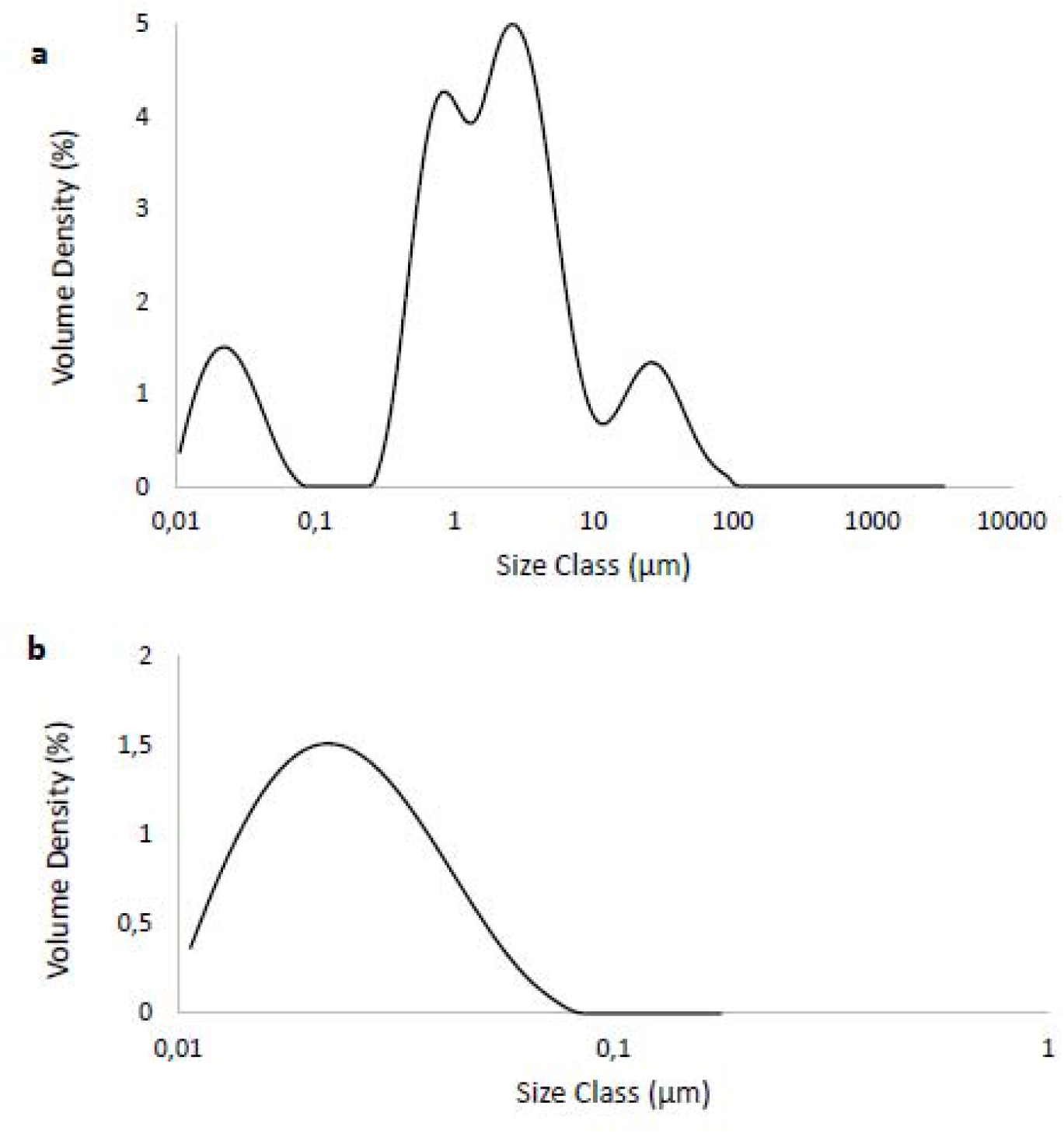
Size of Fe_3_O_4_ Nanoparticles (NP): (a): Involving all particles in the sample; (b) Nanosize cut of the particles

These results show that the sample used also contained particles larger than nanoparticles. The average size, including the two populations, was 180 ± 0.05 nm. This behavior of larger sizes found for Fe_3_O_4_ NP was also reported by Hansein et al. ^21^ who found sizes between 96-400 nm. In the work by Abdsalam et al.^20^, Fe_3_O_4_ NP sizes did not exceed 7 ± 0.2 nm. It is worth mentioning that the NPs used by Abdsalam et al. ^20^ were synthesized, and the NPs used by the present study and by Hansein et al. ^21^ were obtained commercially. The size of NP is extremely important for the process as it can affect the binding and activation of membrane receptors and the expression of proteins^30^, thus acting to stimulate the growth of methanogenic archaea ^31^.

For better visualization, a cut in the graph was made of particles found only in nano size, which are shown in Figure 1b. The average size of these Fe_3_O_4_ NP was 23.56 ± 0.05 nm, which can be considered a greater size as some authors have reported a decrease in CH_4_ production by using Fe NPs greater than 55 nm ^21, 32^. Another important factor is that these Fe_3_O_4_ NPs used in this work have a spherical shape (Figure 2), and this improved CH_4_ production in the work of Abdsalam et al. ^20^ which is explained by the greater membrane wrapping time required for the elongated particles.

**Figure 2.**
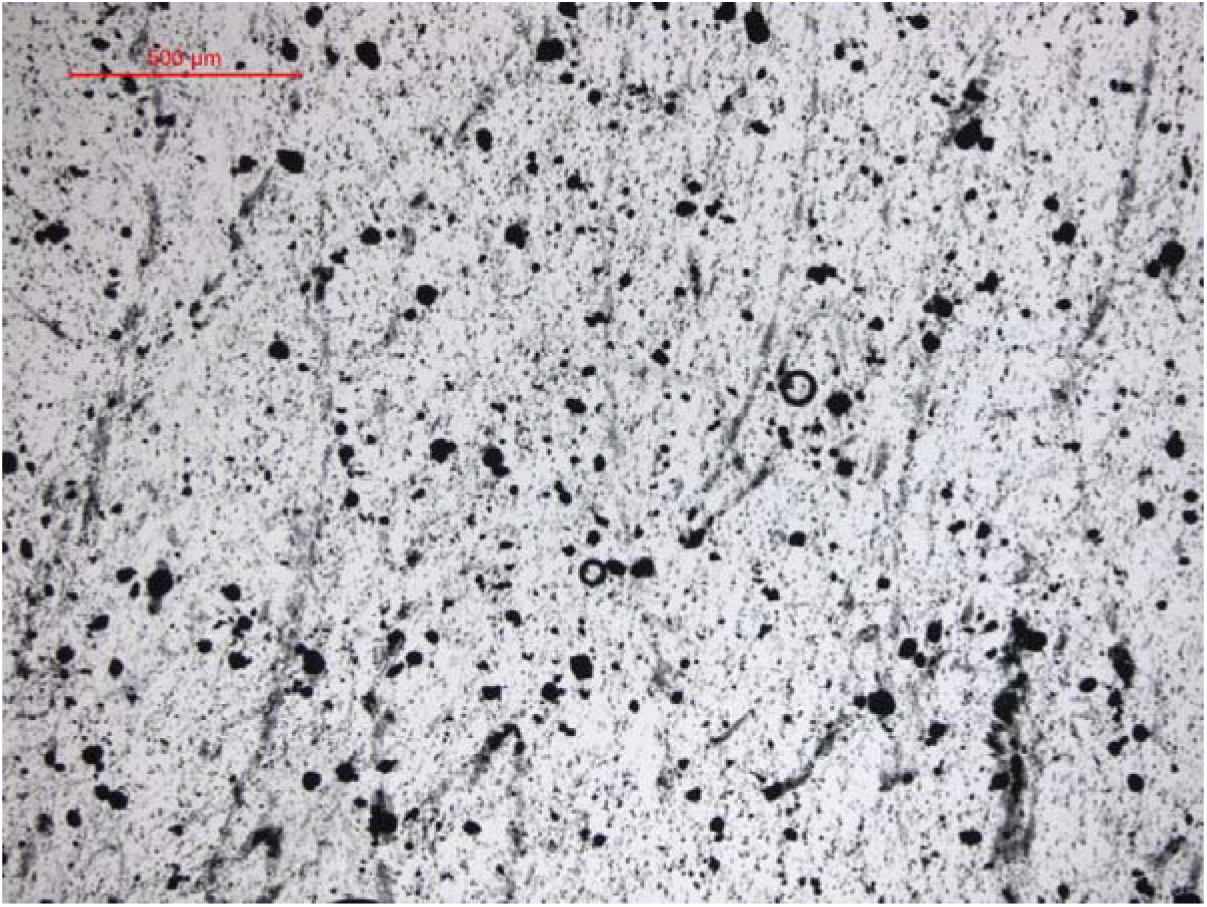
Mastersize images (50x) showing Fe_3_O_4_ NP

Apart from the particle size, other factors that play a role to stimulate CH_4_ production are: the NP concentration added, the type of substrate and the interaction between them ^18^. The preliminary tests of BMP with different concentrations of NP Fe_3_O_4_ enabled us to assess such factors. The zeta potential (ZP) analysis of the nanoparticles was also performed to evaluate their dispersibility in the medium; however, as the NPs used are magnetic particles and have sizes larger than nano, the dispersion remained unstable, which invalidate these results. The same condition has already been reported by Gonzalez et al. ^32^.

### 3.2 Batch preliminary assays

Table 4 shows the results of the specific accumulated CH_4_ in triplicate of each of the tests with different concentrations of Fe_3_O_4_ NP, and Figure 3 presents the cumulative CH_4_ production over time.

**Figure 3.**
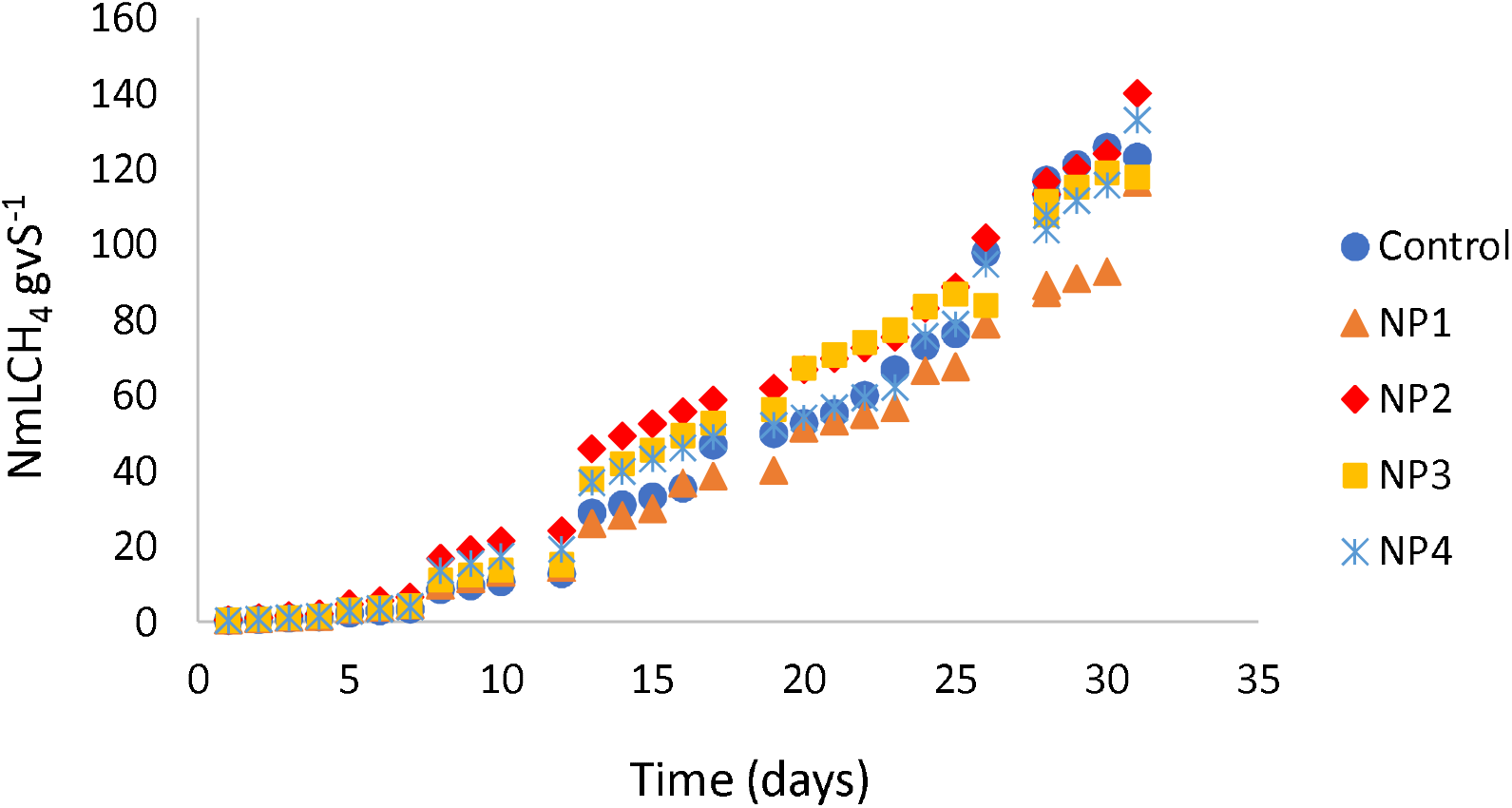
Specific cumulative CH_4_ production according to the concentrations of Fe_3_O_4_nanoparticle (NP) additions to co-digestion. NP1: 1 mg L^-1^; NP2: 5 mg L^-1^; NP3: 10 mg L^-1^; NP4: 20 mg L^-1^

**Table 4.** Final specific cumulative CH_4_ production from the co-digestion in different concentrations of Fe_3_O_4_ NP

| Assay | Cumulative CH<sub>4</sub> (NmLVS<sup>-1</sup>)<sup>a</sup> |
| --- | --- |
| Control | 123.24 ± 9.60 |
| NP 1 | 116.49 ± 17.45 |
| NP 2 | 140.13 ± 95.60 |
| NP 3 | 117.90 ± 10.68 |
| NP 4 | 133.02 ± 106.29 |

<sup>a</sup>: mean of three replicates ± standard variation

There was no significant difference between treatments with different concentrations of Fe_3_O_4_ NP (p-value = 0.1357 with p < 0.05). Although there was no significant difference, NP 2, NP 3 and NP 4 assays presented higher CH_4_ production than the control (Figure 3), while NP 1 was below the control. The NP 2 test showed a 13% increase in CH_4_ production compared to the control, while the NP 4 test showed a 7% increase in CH_4_ production (Table 4).

In the work of Hansein et al. ^21^, a 25% increase in CH_4_ production was obtained, using 15 mg L^-1^ of NP Fe_3_O_4_ in BMP assays, with poultry litter residues, under mesophilic conditions. However, higher CH_4_ concentration (34%) was obtained using NP Ni with a concentration of 12 mg L^-1^. The specific substrate characteristics, the experimental conditions, and the origin of the inoculum may influence these differences in production. The addition of Fe_3_O_4_ NPs may also cause the short lag phase, according to Krongthamchat et al. ^33^.

Although no significant difference in CH_4_ production was observed in the preliminary tests, the concentration of 5 mg L^-1^ of NP Fe_3_O_4_ (NP 2 experiment) was chosen to be applied in the s-CSTR reactor, which showed a higher increase in the CH_4_ production compared to the control. It is also known that nanoparticles are not easy to be separated from biodegradable waste, which may subsequently cause an accumulation of inorganic pollutants (usually heavy metals) inside anaerobic digesters ^34^. For this reason, the selection of the lowest NPs concentration was reinforced to cause less environmental impact on AD. The differences in the operation of the continuous reactor with the addition of NP were compared to the same reactor operation, but without the addition of NP ^8^.

### 3.3 Analysis of s-CSTR operational efficiency

#### 3.3.1 Biogas generations and reactor performance

Figure 4 shows the results obtained from CH_4_ production and organic matter removal according to the different applied OLRs. Along phases I and II, an intense variation in the organic matter removal was observed, from 30% to 70%, while the CH_4_ production remained stable, ranging from 0.1 to 0.5 NLCH_4_ gVS^-1^. These variations are characteristic of the acidogenic phase, marking the reactor start-up. After approximately 60 days (phase III), both the CH_4_ production and the organic matter removal maintained higher stability, indicating that the reactor entered the methanogenic phase. Between phase IV and phase V, the CH_4_ production remained around 0.5 and 1 NLCH_4_ gVS^-1^, within an increase trend. In phase VI, after the addition of Fe_3_O_4_NP, there was a 40% increase in CH_4_ production (122 days), corresponding to the highest CH_4_ production throughout the entire operation: 2.8 ± 0.1 NLCH_4_ gVS^-1^ and removal of 71% ± 0.9% of organic matter. In phase VII, CH_4_ production started to decrease, although the organic matter removal remained stable. In phase VIII, CH_4_ production remained low (0.09 ± 0.03 NLCH_4_ gVS^-1^) and organic matter removal continued to decrease (51 ± 2.8%), reaching the collapse of the reactor in phase IX.

**Figure 4.**
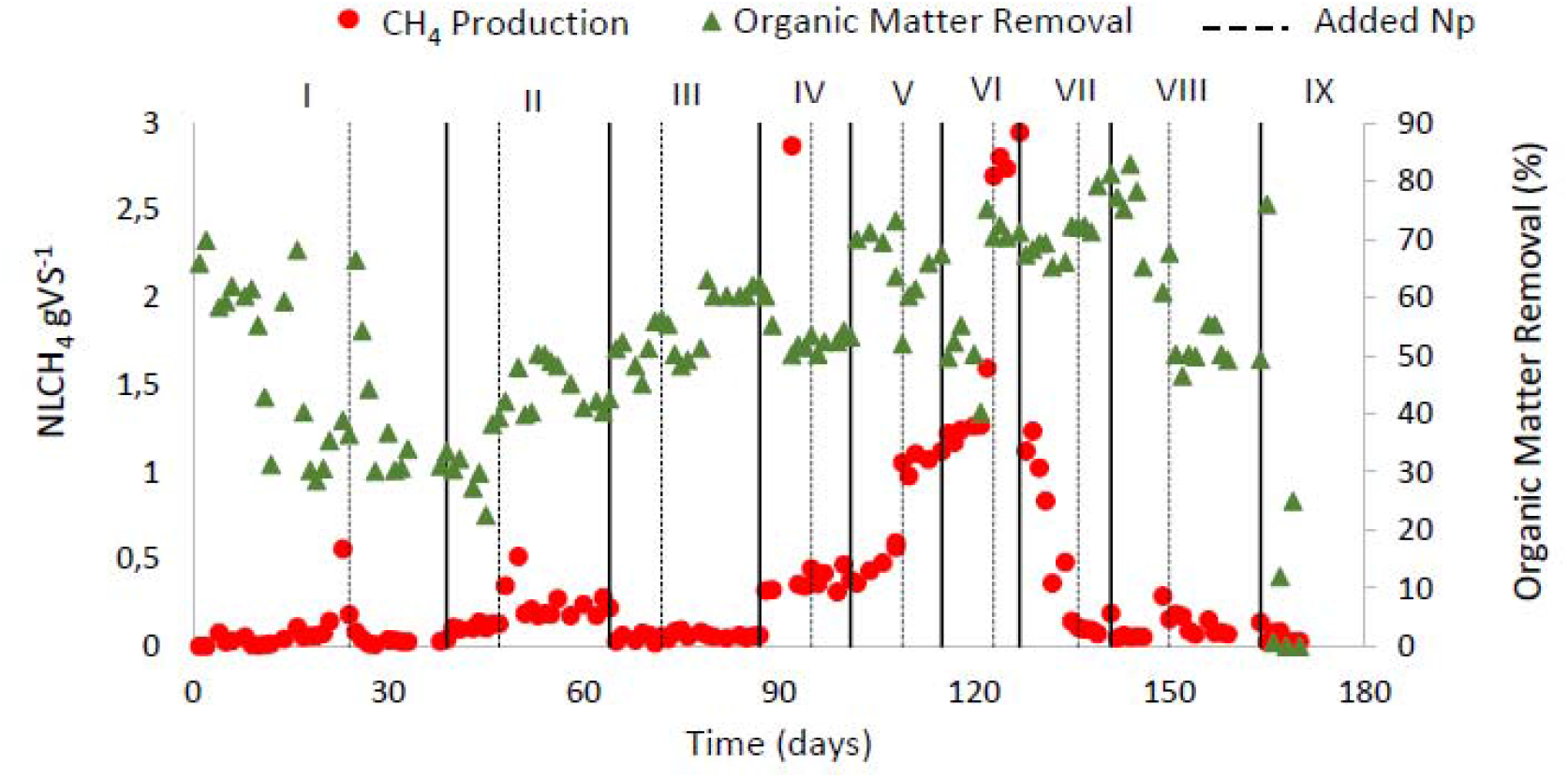
Methane production and organic matter removal with the reactor operation according to the applied OLRs (g VS L^-1^ day^-1^): 2.0 (Phase I); 2.35 (Phase II); 3.0 (Phase III); 4.0 (Phase IV); 4.7 (Phase V); 5.5 (Phase VI); 6.6 (Phase VII); 8.0 (Phase VIII); 9.0 (Phase IX)

In our previous study Volpi et al. ^8^, the co-digestion of the same residues in a semi-CSTR without NP addition reached 0.233 ± 1.83 NLCH_4_ gSV^-1^ and 83.08 ± 13.30% organic matter removal. Compared to the present work, the CH_4_ production from the semi-CSTR containing NP was 91% higher, which confirms that the presence of Fe_3_O_4_ NP contributed to better development and performance of the microbial community concerning the organic matter conversion to CH_4_ as Fe is a growth stimulant of methanogenic Archaea and they are dependent on this element to enzyme synthesis ^14, 35^. In addition, the maximum CH_4_ production and the reactor collapses from the previous ^8^ occurred in the OLRs of 4.16 gVS L^-1^ day^-1^ and 5.23 gVS L^-1^ day^-1^, respectively, while in the present work this occurred in the respective OLR of 5.5 gVS L^-1^ day^-1^ and 9 gVS L^-1^ day^-1^. This fact indicates that the presence of Fe_3_O_4_NP allows the application of larger OLRs, resulting in larger fed volumes to the reactor and, consequently, allowing the treatment of larger waste volumes.

Hassanein et al. ^21^ obtained maximum cumulative production of 339 mLCH_4_ gVS^-1^ from poultry litter in BMP assays, with the addition of 15 mg L^-1^ Fe_3_O_4_ NP, while Abdsalam et al. ^20^ reported 351 mLCH_4_ gVS^-1^ from manure in BMP tests, with 20 mg L^-1^ of Fe_3_O_4_ NP addition. The literature reported using NP in BMP assays and smaller vials to assess NP activity. It is worth mentioning that the use of the substrate with the type of NP interferes with CH_4_ production and NP concentration has also been used to interfere. In the work by Abdsalam et al. ^18^ and Uemura ^36^, it was confirmed that Ni NP was the one that best impacted the increase in CH_4_ production in the use of municipal solid waste. In the study by Ali et al. ^28^, four concentrations of Fe_3_O_4_ NPs (50, 75, 100, and 125 mg L^-1^) were tested in assays with municipal solid waste. The results showed that the addition of 75 mg L^-1^ Fe_3_O_4_NPs increases the CH_4_ production by 53.3%. In contrast, less CH_4_ production was observed by adding a high concentration of Fe_3_O_4_ NPs.

Absalam et al. ^18^ showed that the addition of Fe_3_O_4_ magnetic NPs increased microbial activity during start-up to 40 days of HRT. However, in the present study, an increase in microbial activity was observed in the middle of the operation (phases IV, V, and VI, after 90 days), in agreement with Quing Ni et al ^35^ who indicated that during the exposure of 50 mg L^-1^ of magnetic NPs the adverse effects were insignificant in microbial consortium and concluded that magnetic NPs appeared to be non-toxic during long-term contact. The best performance was due to the presence of Fe2^+^ / Fe3^+^ ions, introduced into the reactor in the form of nanoparticles that could be adsorbed as the growth element of anaerobic microorganisms ^18^. In addition, Fe_3_O_4_ magnetic NP ensures a distribution of the iron ions in the slurry through the corrosion of the NPs, thus maintaining the iron requirement of the reactor supplied ^18^. The presence of NPs also shows a possible effect on the hydrolysis-acidification process, increasing the reduction of the substrate as there were increasing amounts of organic matter removed in phases V, VI and VII, and a subsequent increase in CH_4_ production.

#### 3.3.2 Evaluation of pH, ORP and Alkalinity readings

Figure 5a shows the results obtained from the reactor inlet and outlet pH, as well as the results of oxide-reduction potential (ORP).

**Figure 5.**
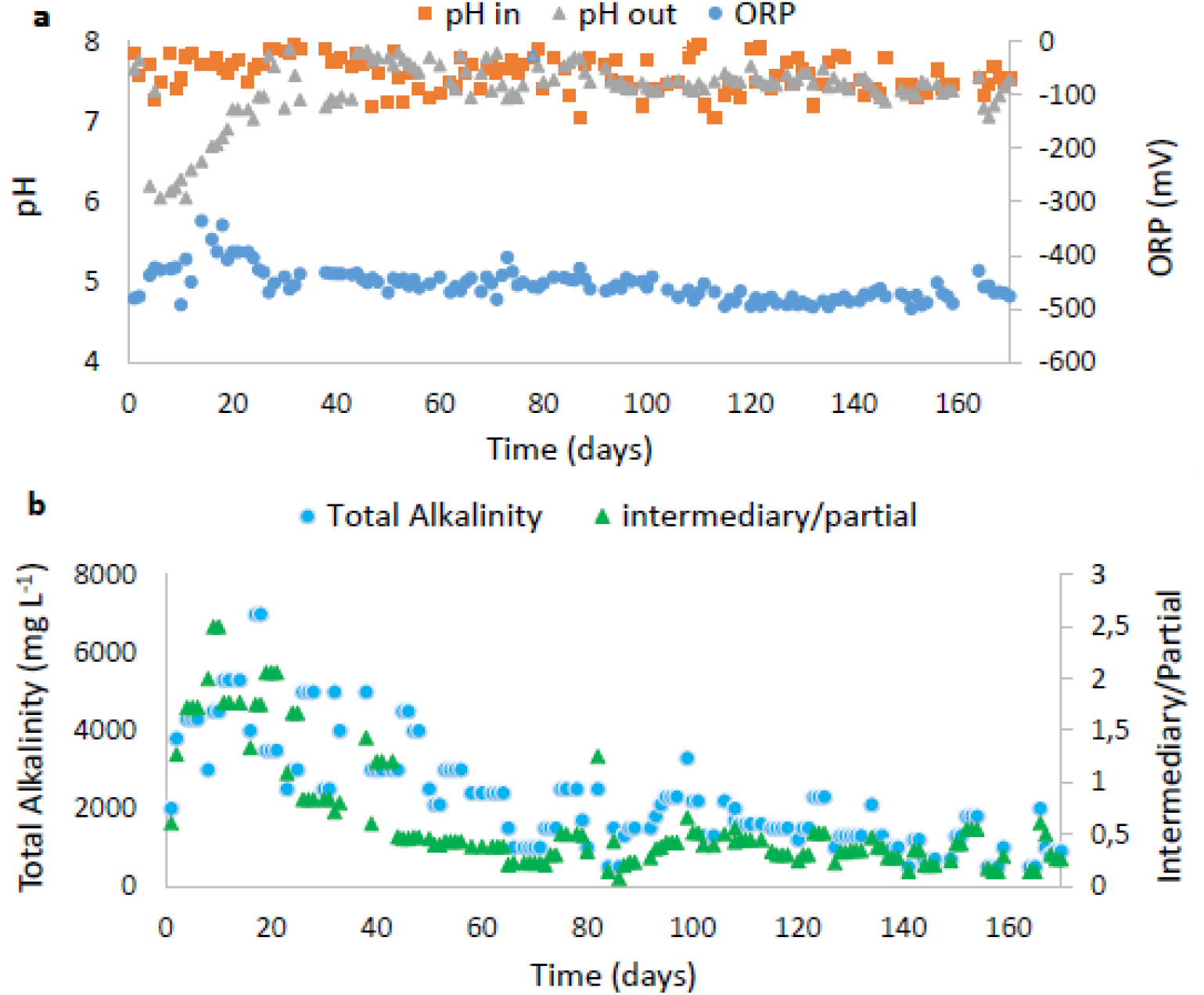
pH, Oxidation Reduction Potential (a) and Alkalinity (b) along with the reactor operation according to the applied OLRs (g VS L^-1^ day^-1^): 2.0 (Phase I); 2.35 (Phase II); 3.0 (Phase III); 4.0 (Phase IV); 4.7 (Phase V); 5.5 (Phase VI); 6.6 (Phase VII); 8.0 (Phase VIII); 9.0 (Phase IX)

The outlet pH in the first days of reactor operation was around 6, and it was necessary to daily adjust the inlet pH to neutrality, while a high variation in the ORP values was detected. These characteristics mark acidogenesis, and the intense oxidation reduction reactions typical of the AD process ^37^. After 60 days, the pH remained between 7.5 and 8 until the end of the operation, indicating that from this date on the reactor entered the methanogenesis phase: the pH remained stable and no more pH adjustment at the reactor inlet was needed. The same occurred for the ORP values, which remained around -460 and -490 mV after 60 days.

In our previous work, methanogenesis was established only after 90 days, with stabilization of the pH and ORP values ^8^. In this present work, methanogenesis was established about 30 days before, indicating that the presence of NPs played a role in this fact, as the addition of Fe_3_O_4_ NP has already been reported to reduce the AD lag phase ^20^. In addition, Feng et al. ^38^ showed that the addition of Fe in the AD system can directly serve as an electron donor to reduce CO_2_ into CH_4_ through autotrophic methanogenesis causing improvement of CH_4_ production, according to reactions (1), (2) and (3).

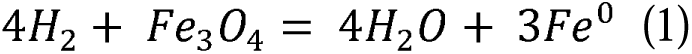

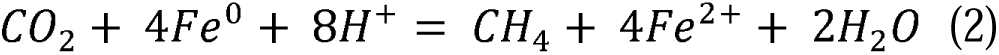

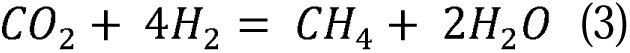

From this process, the substrates are deprived of hydrogen ions (H^+^) which increases the pH of the substrate, and the capture of CO_2_ also prevents the formation of carbonic acid, increasing the pH of the substrate ^20^. This may explain the increase in pH after 24 days (Table 3) as it was the first addition of Fe_3_O_4_ NPs. The methanogenesis process was also stimulated, as this nano additive served as an electron donor that could reduce CO_2_ into CH_4_.

At the beginning of the operation, the ORP varied between -350 and -550 mV, and this variation is a characteristic of acidogenesis and reactor start-up ^8^. However, this variation in a start-up was much smaller than that reported in Volpi et al. ^8^ (−800 and −300 mV) indicating greater stability of the operation. After approximately 40 days (Figure 5a), it was observed that the ORP remained practically constant until the end of the operation, varying between -480 and -400 mV, although the literature shows that the ORP values characteristic from the acidogenic and methanogenic phase are between - 330 and -428 mV ^39^. These low ORP values in the system are characteristic of the presence of Fe NPs as they reduce the system’s ORP to increase the conversion of complex compounds to volatile fatty acids and to be able to provide ferrous ions for the growth of fermentative and methanogenic Archeae ^40^.

It is important to demonstrate that the practically constant ORP values are in agreement with the OA values (Section 3.3.3), which are in extremely low concentrations when the reactor stabilizes in methanogenesis. Here it is worth emphasizing that the differences in the ORP values found in the literature are varied due to the different raw materials applied, experimental conditions and the type of NP used.

Figure 5b shows the results of alkalinity obtained during the operation. It can be observed that the alkalinity was high up to 60 days, and followed the presence of OA (Figure 6a, Section 3.3.3) characterizing the acidogenic step of the process. After 60 days, the intermediate/partial alkalinity (IA/PA) was below 0.3, which is considered ideal for AD, as it demonstrates stability ^41^. Similar to the behavior of the ORP, the IA/PA also remained stable throughout the process, showing self-regulation of methanogenesis. In our previous study ^8^, such alkalinity stability only happened after 90 days, confirming the hypothesis that the presence of Fe_3_O_4_NP has reduced the lag phase. In addition, Fe3O4NP can absorb inhibitory compounds and act as a pH buffer, further improving the alkalinity of the process.

**Figure 6.**
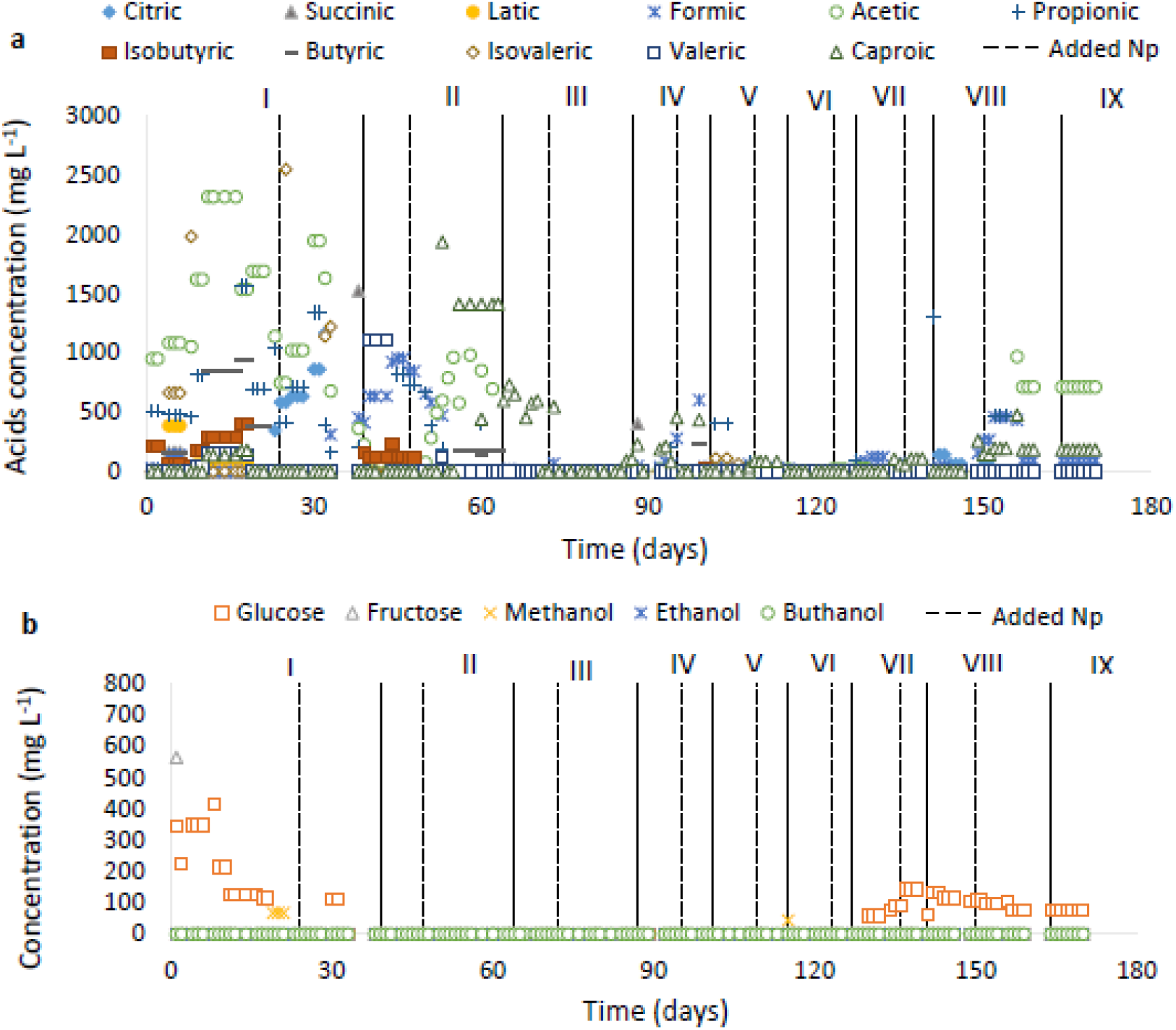
Values of Organic acids (a), Carbohydrates, and Alcohols (b) along with the reactor operation based on to the applied OLRs (g VS L^-1^ day^-1^): 2.0 (Phase I); 2.35 (Phase II); 3.0 (Phase III); 4.0 (Phase IV); 4.7 (Phase V); 5.5 (Phase VI); 6.6 (Phase VII); 8.0 (Phase VIII); 9.0 (Phase IX)

#### 3.3.3 Assessment of degradation pathways: OA, Carbohydrate and Alcohol indications

Figure 6 shows the results obtained from OA and carbohydrates and alcohols.

In phases I and II (Figure 6a), the presence of high concentrations of OA was in accordance with the reactor start-up (the acidogenic phase). After 60 days, the concentrations of these OA considerably decreased, indicating the entrance of the reactor in the methanogenic phase and agreeing with what was discussed in the previous sections (Sections 3.3.1, 3.3.2).

At the beginning of the reactor operation (phase I), the concentration of acetic acid was relatively high, which is favorable for the CH_4_ production process as it is the main precursor of the CH_4_ metabolic route ^42^. In addition to acetic acid, there was also the presence of propionic acid, which can be inhibitory to the metabolic pathway of CH_4_ production in concentrations above 1500 mg L^-1^ ^43^. However, this concentration decreased in phase I and phase II, and the acetic acid concentration increased at the end of phase II, indicating that the route of conversion of propionic acid to acetic acid may have prevailed at the beginning of the operation, as also occurred in our previous study^8^. It is worth mentioning that in the presence of low H_2_ pressure, propionic acid consumption is favored ^42^, and Fe is a trace element whose main substrate for oxidation reduction reactions is H_2_ ^14^. The presence of Fe_3_O_4_ NP may have favored the consumption of H_2_, according to reaction (4) and reaction (1), and consequently contributed to the consumption of propionic acid, favoring the formation of acetic acid and this having been converted to CH_4._

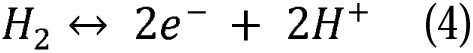

In phases I and II, the presence of formic acid can be observed, and its conversion to acetic acid is typical of acidogenesis ^14^. Therefore, in addition to the conversion of propionic acid to acetic acid, the conversion of formic acid to acetic acid may also have occurred at the end of acidogenesis, marking the beginning of methanogenesis (phase III-Figure 6a). In addition, the presence of Fe NP can increase the production of acetate and donate electrons for direct conversion of CO_2_ into CH_4_ by autotrophic via methanogenesis ^38^.

The Fe (III) reduction reaction is a favorable process to directly oxidize organics into simple compounds ^44^, increasing the consumption of OA, and eliminate compounds that may be toxic to the process, by stimulating microbial growth, synthesis of necessary enzymes within the oxidation-reduction reactions and consequently greater efficiency in the digestion of organic matter ^14, 40^. The positive effect of Fe (III) supplementation was attributed to the favorable redox conditions, which all avoided the thermodynamic limitations on organic acid degradation. Furthermore, Fe (III) can precipitate H_2_S minimizing related inhibition phenomena ^44^. The control of OA can allow a greater capacity of feed of the digester, without affecting the performance of digestion significantly ^45^, this is what happened in the present study as the OLRs used were higher than the experiment previous ^8^, with higher fed volumes, and a stable operation, reaching higher CH_4_ production.

The presence of caproic acid draws attention at the end of phase II and the beginning of phase III (Figure 6a). Caproic acid is produced by lengthening the chain of short-chain volatile fatty acids, such as acetic acid and butyric acid through an oxidation reaction, in which some species gain energy by increasing the length of the volatile organic acids chain with reducing substrates such as ethanol and lactic acid ^46^. However, in the operation, neither the presence of lactic acid nor ethanol was detected (Figure 6a and Figure 6b), but it seems that Fe_3_O_4_NP may have acted as this reducing substrate, gaining electrons and allowing an increase in the chain of butyric and acetic acids. This fact may also have been caused by the continuous feeding process of the reactor, in which Fe_3_O_4_NP was added with a certain frequency, having a constant availability of the electron donor for the formation of caproic acid and in agreement with what was reported by Owusu-Agyeman et al. ^46^. Even with the possible change of the route for caproic acid production, CH_4_ production prevailed, indicating the self-regulation of the microbial consortium for the metabolic route of CH_4_. Although not the focus of this work, the addition of Fe_3_O_4_ NP with the residues of the sugarcane industry can stimulate the caproic acid production, an organic acid that has high added value because it is used as antimicrobials for animal feed and precursor aviation fuel ^47^.

Figure 6b shows that at the beginning of the operation, there was greater availability of glucose, and when the reactor entered the methanogenesis phase, the glucose concentration was very low, indicating the self-regulation of the process for CH_4_ production. When the reactor began to decrease its CH_4_ production, phases VII and VIII, the concentration of glucose increased again, indicating the start of the reactor collapse.

### 2.6 Assessing the diversity of microbial communities

Figure 7 shows the observed values of richness (number of species-a), and the calculated values from diversity (Shannon index-b) and wealth estimate (Chao1 estimator-c) of the samples. The results show that the number of species (Figure 7a) and richness (Figure 7b) of the A1 samples was higher than that of the A2 sample. This behavior is as expected for these results as the A1 samples are samples from the initial inoculum, that is, from the inoculum without having been inserted into the reactor. The A2 samples are from the inoculum when the CH_4_ production was stabilized, that is to say, that the microbial community present is already “selected” for the specific metabolic route of CH_4_ production according to the substrates used. In addition, the inoculum of Sample A1 comes from a mesophilic reactor, while Sample A2 comes from a thermophilic reactor. Process temperature differences may also have led to this difference between species of microorganisms. These results are consistent with our previous work ^8^ indicating that the presence of NP did not influence the diversity of microorganisms and the change in the microbial community from one sample to another.

**Figure 7.**
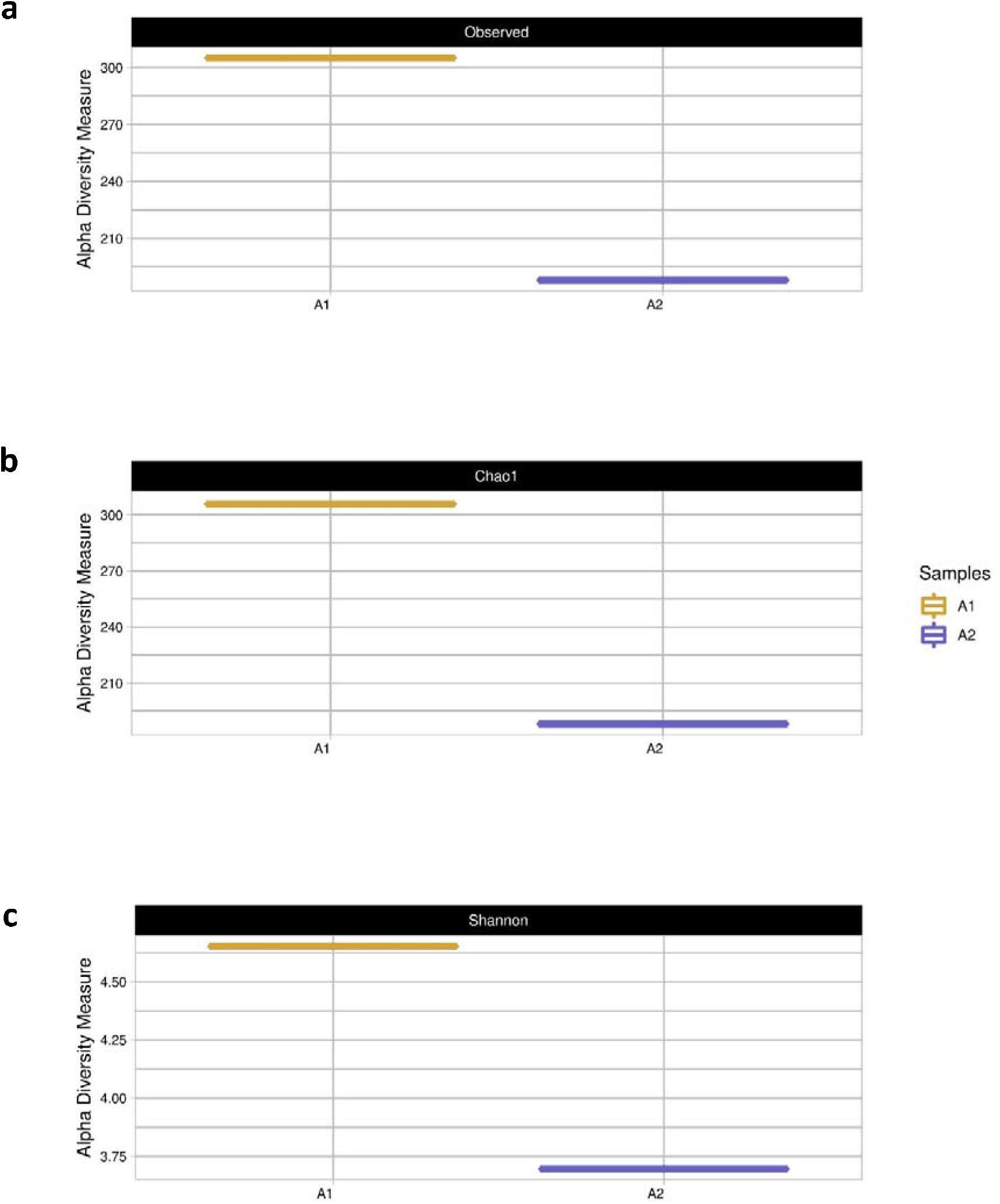
The recorded values of (A) species abundance (number of species), (B) estimated species richness (Chao1 estimator), and (C) computed diversity values (Shannon index) for Sample 1 (seed sludge) and Sample 2 (sludge from Phase IV of stable s-CSTR operation).

The Shannon index obtained from sample A1 was close to 5.0, while that from sample A2 was less than 3.75. As discussed in ^8^, when the value of the Shannon index is greater than 5.0, it indicates greater microbial diversity in anaerobic digesters ^48^. Thus, it can be seen that the A2 sample has a much lower microbial diversity than the A1 as these microorganisms are in stabilized metabolic routes for CH_4_ production, indicating that this microbial community is even more specific.

Figure 8 shows the results obtained from phylum in relation to *Bacteria* order (a) and *Archaea* order (b) from samples A1 and A2.

**Figure 8.**
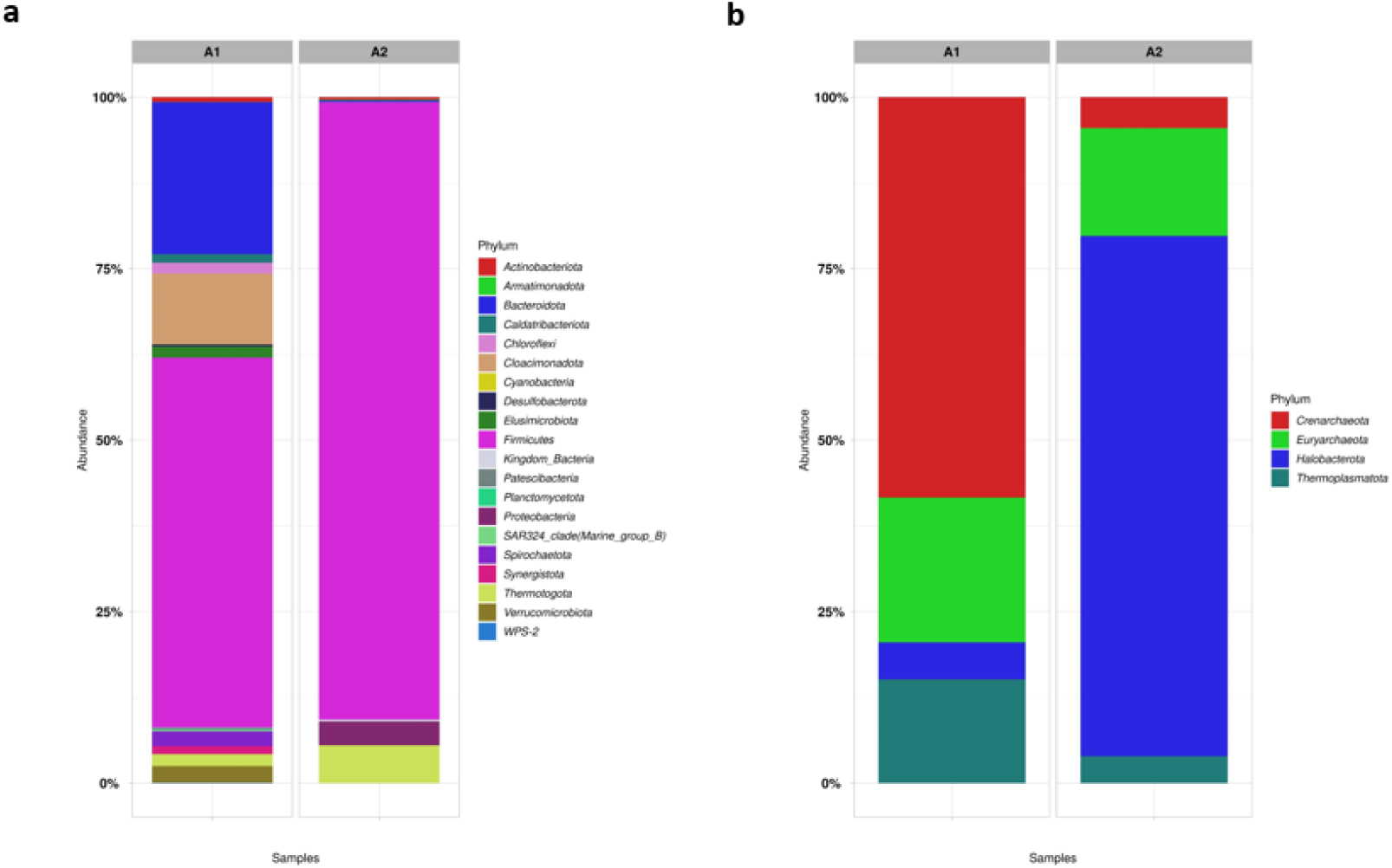
The proportional representation of microorganisms at the phylum level in the Bacteria order (a) and Archaea order (b) in the seed sludge (Sample A1) and the s-CSTR sludge with stable CH_4_ production (Sample A2).

Following what was discussed above, the phyla variety found in sample A1 (Figure 8a) was much larger than those found in A2. In sample A1, the principal phyla found from *Bacteria* order was: (∼25%) *Bacteroidota*, (∼15%) *Cloacimonadota*, (∼50%) *Firmicutes* and (∼2%) *Spirochaetota*. Microorganisms of the phylum *Bacteroidota, Cloacimonadota,* and *Spirochaetota* are generally found in mesophilic processes and are bacteria responsible for the fermentative and hydrolytic steps of AD ^49, 50^. The presence of these three phyla in the A1 sample and the absence of them in the A2 sample indicates how the temperature influenced the change in the bacterial community as the A1 inoculum comes from a mesophilic process. The large presence of the *Firmicutes* phylum is to be expected as they are one of the main phyla of anaerobic processes, and most cellulolytic bacteria belong to them ^51^. In sample A2, the main phyla found are (∼80%) *Firmicutes*, (∼2%) *Protobacteria,* and (∼5%) *Thermotogota*. The *Thermotogota* phylum is characteristic of thermophilic processes ^52^, and bacteria of the *Protobacteria* phylum are characteristic for degrading lignocellulosic material ^51^. It is important to mention that these two last phyla are present in smaller proportions in sample A1, indicating the possibility of a change in the microbial community due to experimental conditions and used substrates. Furthermore, in our previous work ^8^, these same phyla were found in the sample when the reactor was stabilized for CH_4_ production, indicating that the presence of Fe_3_O_4_ NP did not influence the change in the microbial community concerning Bacteria order. Zhang et al. ^50^showed that the presence of *Proteobacteria* followed by *Firmicutes* were the central syntrophic acetogenins for propionate oxidation via the methylmalonyl-CoA pathway, perhaps indicating the presence of this metabolic route when CH_4_ production stabilized, as discussed in Section 3.3.3.

Concerning *Archaea* order phyla, in sample A1, *Euryarchaeota* was observed (∼25%) and in sample A2 (∼20%) of the same phylum. This phylum is characteristic of methanogenic *Archaeas*, responsible for CH_4_ production. In addition to this main phylum, other phyla of the *Archaea* order were also found, such as *Crenarchaeota, Halobacterota*, but they are not highly relevant for the results of CH_4_ production, which was the focus of this work.

Figure 9 shows the main genera found for samples A1 and A2 to the Bacteria order (a) and the Archaea order (b). As previously discussed, the A1 sample presented a very large microbial diversity, with no genus that was predominant in the process concerning the *Bacteria* order. Its genera of microorganisms come from the main phyla (*Bacteroidota, Cloacimonadota, Firmicutes*) and are characteristic of acidogenic and hydrolytic processes.

**Figure 9.**
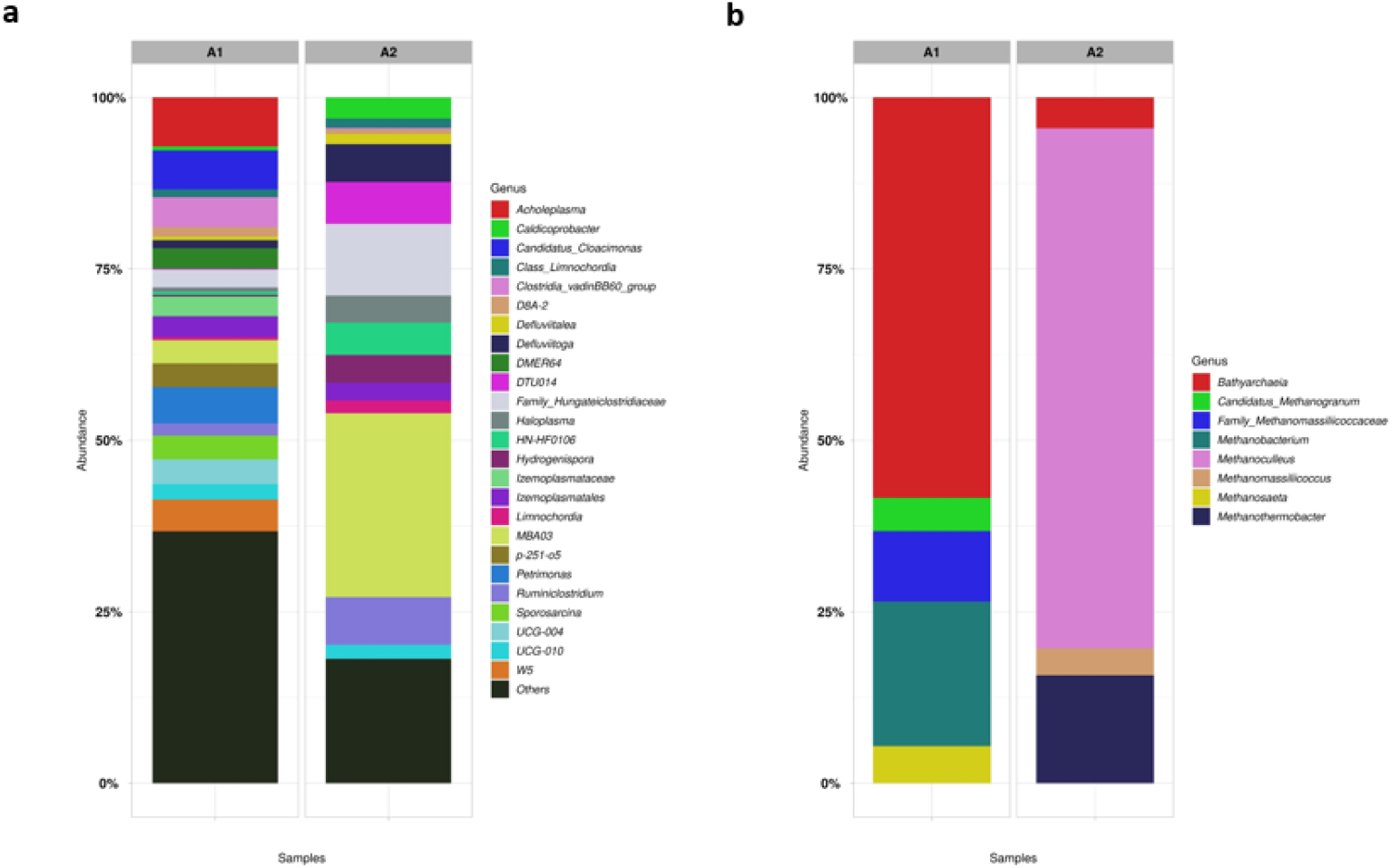
The proportional representation of microorganisms at the genus level in the Bacteria order (a) and Archaea order (b) in the seed sludge (Sample A1) and the s-CSTR sludge with stable CH_4_ production (Sample A2).

Sample A2 has some genera of the Bacteria order that stand out, such as (∼5%) *Defluvitoga*, (∼3%) *Hydrogenispora*, (∼9%) *Ruminiclostridium*. These genera were also present in the reactor operation without the presence of Fe_3_O_4_ NP ^8^.

*Defluvitoga* genus, belonging to the phylum *Thermotogota*, is reported to be dominant in the degradation of organic materials in CSTRs or thermophilic bioelectrochemical reactors ^53^. *Ruminiclostridium*, belonging to the phylum *Firmicutes*, are hydrolytic bacteria characterized by metabolizing cellulosic materials, with a high concentration of lignocellulose ^54^, which is the case of residues used in reactor operation. In the work by Kang et al. ^55^, wheat straw was used for AD, and bacteria belonging to the genus *Ruminiclostrium* and *Hydrogenispora* were found as the main microorganisms. This fact leads to the association that such bacteria are present in the degradation of lignocellulose substrates as wheat straw and residues from the present work have a similar composition.

*Hydrogenispora* is acetogenic bacteria, which can ferment carbohydrates such as glucose, maltose, and fructose into acetate, ethanol, and H_2_ ^55^. These bacteria can act in conjunction with hydrogenotrophic methanogens. In Figure 9b, the predominant methanogenic *Archaea* in sample A2 was (∼70%) *Methanoculleus*. This methanogenic *Archaea* is characterized by acting syntrophic oxidation of acetate (SAO) coupled with hydrogenotrophic methanogenesis ^56^. Furthermore, it was also the main methanogenic found in the work by Volpi et al. ^8^. Therefore, despite the addition of NP in the reactor, the presence of the microbial community was not altered and therefore, the metabolic routes were also the same. The presence of NPs only encouraged the activity of the methanogenic *Archaea*, since the substrates used and experimental conditions were the same, there was no change in the metabolic route. The genus (∼15%) *Methanotermobacter* was also found in sample A2. This genus is characterized by being present in thermophilic AD and belongs to the obligate-hydrogenotrophic methanogens^57^. This fact corroborates the possibility that the predominant metabolic route in the co-digestion of vinasse, filter cake, and deacetylation liquor is acetate (SAO) coupled with hydrogenotrophic methanogenesis. Furthermore, it was discussed in Section 3.3.3 that in the presence of low H_2_ pressure, propionic acid consumption is favored, and Fe is a trace element whose main substrate for oxidation-reduction reactions is H_2_. This confirms the fact that the presence of Fe_3_O_4_ NP may have reinforced that the main metabolic pathway for the co-digestion of these residues is through hydrogenotrophic methanogens.

In sample A1 (Figure 9a), (∼20%) *Methanobacterium* and (∼7%) *Methanosaeta* were found*. Methanobacterium* is known as hydrogenotrophic methanogens while *Methanosaeta* is known as obligate-acetoclastic methanogen and has a strong affinity to acetate ^57^. These two genera are not found in sample A2, indicating there was a change in the microbial community from sample A1 to A2 due to different substrates and experimental conditions.

## 4. CONCLUSIONS

The use of Fe_3_O_4_ NP allowed to optimize the biological process of co-digestion of 1G2G ethanol industry residues, providing an increase of approximately 90% in CH_4_ production. The concentration of 5 mg L^-1^ of Fe_3_O_4_ NP was ideal for a stable continuous operation, with production stimulation, without process inhibitions.

These nanoparticles proved to favor the reduction of the lag phase of the process, through a stabilized reactor operation. The reactor collapsed in OLRs of 9 gVS L^-1^ day^-1^, which was an ORL almost 2 times larger than that used in the operation without the presence of NP (9 *vs* 5 gVS L^-1^ day^-1^). Furthermore, the methanogenesis was stabilized after 60 days of operation, which was 30 days earlier than the operation without the addition of NP.

Fe_3_O_4_ NP did not influence the possible metabolic pathways of the process; on the contrary, they stimulated the growth of methanogenic *Archaea*, reinforcing that the main metabolic pathway of these residues in co-digestion is through hydrogenotrophic methanogenesis. *Methanoculleus* were the main methanogenic *Archea* found in the process, and *Defluvitoga, Ruminiclostridium,* and *Hydrogenispora* were the main genus of the *Bacteria* order in process, both with or without the addition of NP.

## REFERENCES

(1) Deublin, D.; Steinhauser, A. Biogas from Waste and Renewable Resources: An Introduction; Wily Online Library: Weinheim, Germany, 2008. https://doi.org/1010029783527621705.

(2) Djalma Nunes Ferraz Júnior, A.; Koyama, M. H.; de Araújo Júnior, M. M.; Zaiat, M. Thermophilic Anaerobic Digestion of Raw Sugarcane Vinasse. Renew Energy 2016. https://doi.org/10.1016/j.renene.2015.11.064.

(3) Fuess, L. T.; Kiyuna, L. S. M.; Ferraz, A. D. N.; Persinoti, G. F.; Squina, F. M.; Garcia, M. L.; Zaiat, M. Thermophilic Two-Phase Anaerobic Digestion Using an Innovative Fixed-Bed Reactor for Enhanced Organic Matter Removal and Bioenergy Recovery from Sugarcane Vinasse. Appl Energy 2017, 189, 480–491. https://doi.org/10.1016/j.apenergy.2016.12.071.

(4) Moraes, B. S.; Zaiat, M.; Bonomi, A. Anaerobic Digestion of Vinasse from Sugarcane Ethanol Production in Brazil: Challenges and Perspectives. Renewable and Sustainable Energy Reviews 2015, 44, 888–903. https://doi.org/10.1016/j.rser.2015.01.023.

(5) Hagos, K.; Zong, J.; Li, D.; Liu, C.; Lu, X. Anaerobic Co-Digestion Process for Biogas Production: Progress, Challenges and Perspectives. Renewable and Sustainable Energy Reviews 2017, 76 (November 2016), 1485–1496. https://doi.org/10.1016/j.rser.2016.11.184.

(6) Janke, L.; Leite, A.; Nikolausz, M.; Schmidt, T.; Liebetrau, J.; Nelles, M.; Stinner, W. Biogas Production from Sugarcane Waste: Assessment on Kinetic Challenges for Process Designing. Int J Mol Sci 2015, 16 (9), 20685–20703. https://doi.org/10.3390/ijms160920685.

(7) Volpi, M. P. C.; Brenelli, L. B.; Mockaitis, G.; Rabelo, S. C.; Franco, T. T.; Moraes, B. S. Use of Lignocellulosic Residue from Second-Generation Ethanol Production to Enhance Methane Production through Co-Digestion. Bioenergy Res 2021. https://doi.org/10.1007/s12155-021-10293-1.

(8) Volpi, M. P. C.; Junior, A. D. N. F.; Franco, T. T.; Moraes, B. S. Operational and Biochemical Aspects of Co-Digestion (Co-AD) from Sugarcane Vinasse, Filter Cake, and Deacetylation Liquor. Appl Microbiol Biotechnol 2021. https://doi.org/10.1007/s00253-021-11635-x.

(9) Moraes, B. S.; Junqueira, T. L.; Pavanello, L. G.; Cavalett, O.; Mantelatto, P. E.; Bonomi, A.; Zaiat, M. Anaerobic Digestion of Vinasse from Sugarcane Biorefineries in Brazil from Energy, Environmental, and Economic Perspectives: Profit or Expense? Appl Energy 2014, 113, 825–835. https://doi.org/10.1016/j.apenergy.2013.07.018.

(10) Brenelli, L. B.; Figueiredo, F. L.; Damasio, A.; Franco, T. T.; Rabelo, S. C. An Integrated Approach to Obtain Xylo-Oligosaccharides from Sugarcane Straw: From Lab to Pilot Scale. Bioresour Technol 2020, 123637. https://doi.org/10.1016/j.biortech.2020.123637.

(11) Demirel, B.; Scherer, P. The Roles of Acetotrophic and Hydrogenotrophic Methanogens during Anaerobic Conversion of Biomass to Methane: A Review. Rev Environ Sci Biotechnol 2008, 7 (2), 173–190. https://doi.org/10.1007/s11157-008-9131-1.

(12) Scherer, P.; Lippert, H.; Wolff, G. Composition of the Major Elements and Trace Elements of 10 Methanogenic Bacteria Determined by Inductively Coupled Plasma Emission Spectrometry. Biol Trace Elem Res 1983, 5 (3), 149–163. https://doi.org/10.1007/BF02916619.

(13) Zhang, Y.; Zhang, Z.; Suzuki, K.; Maekawa, T. Uptake and Mass Balance of Trace Metals for Methane Producing Bacteria. Biomass Bioenergy 2003, 25 (4), 427–433. https://doi.org/10.1016/S0961-9534(03)00012-6.

(14) Choong, Y. Y.; Norli, I.; Abdullah, A. Z.; Yhaya, M. F. Impacts of Trace Element Supplementation on the Performance of Anaerobic Digestion Process: A Critical Review. Bioresour Technol 2016, 209, 369–379. https://doi.org/10.1016/j.biortech.2016.03.028.

(15) Yu, B.; Lou, Z.; Zhang, D.; Shan, A.; Yuan, H.; Zhu, N.; Zhang, K. Variations of Organic Matters and Microbial Community in Thermophilic Anaerobic Digestion of Waste Activated Sludge with the Addition of Ferric Salts. Bioresour Technol 2015, 179, 291– 298. https://doi.org/10.1016/j.biortech.2014.12.011.

(16) Demirel, B.; Scherer, P. Trace Element Requirements of Agricultural Biogas Digesters during Biological Conversion of Renewable Biomass to Methane. Biomass Bioenergy 2011, 35 (3), 992–998. https://doi.org/10.1016/j.biombioe.2010.12.022.

(17) Zhang, Y.; Jing, Y.; Zhang, J.; Sun, L.; Quan, X. Performance of a ZVI-UASB Reactor for Azo Dye Wastewater Treatment. Journal of Chemical Technology and Biotechnology 2011, 86 (2), 199–204. https://doi.org/10.1002/jctb.2485.

(18) Abdelsalam, E.; Samer, M.; Attia, Y. A.; Abdel-Hadi, M. A.; Hassan, H. E.; Badr, Y. Comparison of Nanoparticles Effects on Biogas and Methane Production from Anaerobic Digestion of Cattle Dung Slurry. Renew Energy 2016, 87, 592–598. https://doi.org/10.1016/j.renene.2015.10.053.

(19) Abdelsalam, E.; Samer, M.; Attia, Y. A.; Abdel-Hadi, M. A.; Hassan, H. E.; Badr, Y. Effects of Co and Ni Nanoparticles on Biogas and Methane Production from Anaerobic Digestion of Slurry. Energy Convers Manag 2017, 141, 108–119. https://doi.org/10.1016/j.enconman.2016.05.051.

(20) Abdelsalam, E.; Samer, M.; Attia, Y. A.; Abdel-Hadi, M. A.; Hassan, H. E.; Badr, Y. Influence of Zero Valent Iron Nanoparticles and Magnetic Iron Oxide Nanoparticles on Biogas and Methane Production from Anaerobic Digestion of Manure. Energy 2017, 120, 842–853. https://doi.org/10.1016/j.energy.2016.11.137.

(21) Hassanein, A.; Lansing, S.; Tikekar, R. Bioresource Technology Impact of Metal Nanoparticles on Biogas Production from Poultry Litter. Bioresour Technol 2019, 275 (December 2018), 200–206. https://doi.org/10.1016/j.biortech.2018.12.048.

(22) Mu, H.; Chen, Y.; Xiao, N. Bioresource Technology Effects of Metal Oxide Nanoparticles (TiO 2, Al 2 O 3, SiO 2 and ZnO) on Waste Activated Sludge Anaerobic Digestion. Bioresour Technol 2011, 102 (22), 10305–10311. https://doi.org/10.1016/j.biortech.2011.08.100.

(23) Wang, T.; Zhang, D.; Dai, L.; Chen, Y.; Dai, X. Effects of Metal Nanoparticles on Methane Production from Waste-Activated Sludge and Microorganism Community Shift in Anaerobic Granular Sludge. Sci Rep 2016, 6 (April), 1–10. https://doi.org/10.1038/srep25857.

(24) Mamani, J. B.; Gamarra, L. F. Synthesis and Characterization of Fe 3 O 4 Nanoparticles with Perspectives in Biomedical Applications. 2014, 17 (3), 542–549.

(25) APHA, AWWA, W. Standard Methods for the Examination of Water and Wastewater, twenty-sec.; Washington, DC., 2012.

(26) Triolo, J. M.; Pedersen, L.; Qu, H.; Sommer, S. G. Biochemical Methane Potential and Anaerobic Biodegradability of Non-Herbaceous and Herbaceous Phytomass in Biogas Production. Bioresour Technol 2012, 125, 226–232. https://doi.org/10.1016/j.biortech.2012.08.079.

(27) VDI 4630. Fermentation of Organic Materials. Characterization of the Substrate, Sampling, Collection of Material Data, Fermentation Tests. Düsseldorf: Verein Deutscher Ingenieure; 2006.

(28) Ali, A.; Mahar, R. B.; Soomro, R. A.; Sherazi, S. T. H. Fe3O4 Nanoparticles Facilitated Anaerobic Digestion of Organic Fraction of Municipal Solid Waste for Enhancement of Methane Production. Energy Sources, Part A: Recovery, Utilization and Environmental Effects 2017, 39 (16), 1815–1822. https://doi.org/10.1080/15567036.2017.1384866.

(29) Zhang, Z.; Guo, L.; Wang, Y.; Zhao, Y.; She, Z.; Gao, M.; Guo, Y. Application of Iron Oxide (Fe3O4) Nanoparticles during the Two-Stage Anaerobic Digestion with Waste Sludge: Impact on the Biogas Production and the Substrate Metabolism. Renew Energy 2020, 146, 2724–2735. https://doi.org/10.1016/j.renene.2019.08.078.

(30) Jiang, W. E. N.; Kim, B. Y. S.; Rutka, J. T.; Chan, W. C. W. Nanoparticle-Mediated Cellular Response Is Size-Dependent. 2008, 145–150. https://doi.org/10.1038/nnano.2008.30.

(31) Mu, H.; Chen, Y.; Xiao, N. Effects of Metal Oxide Nanoparticles (TiO 2, Al 2O 3, SiO 2 and ZnO) on Waste Activated Sludge Anaerobic Digestion. Bioresour Technol 2011, 102 (22), 10305–10311. https://doi.org/10.1016/j.biortech.2011.08.100.

(32) Gonzalez-estrella, J.; Sierra-alvarez, R.; Field, J. A. Toxicity Assessment of Inorganic Nanoparticles to Acetoclastic and Hydrogenotrophic Methanogenic Activity in Anaerobic Granular Sludge. J Hazard Mater 2013, 260, 278–285. https://doi.org/10.1016/j.jhazmat.2013.05.029.

(33) Krongthamchat, K.; Riffat, R.; Dararat, S. Effect of Trace Metals on Halophilic and Mixed Cultures in Anaerobic Treatment. International Journal of Environmental Science and Technology 2006, 3 (2), 103–112. https://doi.org/10.1007/BF03325913.

(34) Zhu, X.; Blanco, E.; Bhatti, M.; Borrion, A. Impact of Metallic Nanoparticles on Anaerobic Digestion: A Systematic Review. Science of the Total Environment 2021, 757. https://doi.org/10.1016/j.scitotenv.2020.143747.

(35) Ni, S. Q.; Ni, J.; Yang, N.; Wang, J. Effect of Magnetic Nanoparticles on the Performance of Activated Sludge Treatment System. Bioresour Technol 2013, 143, 555–561. https://doi.org/10.1016/j.biortech.2013.06.041.

(36) Sh, U. Mineral Requirements for Mesophilic and Thermophilic Archive of SID. Archive of SID 2010, 4 (1), 33–40.

(37) Vongvichiankul, C.; Deebao, J.; Khongnakorn, W. Relationship between PH, Oxidation Reduction Potential (ORP) and Biogas Production in Mesophilic Screw Anaerobic Digester. Energy Procedia 2017, 138, 877–882. https://doi.org/10.1016/j.egypro.2017.10.113.

(38) Feng, Y.; Zhang, Y.; Quan, X.; Chen, S. Enhanced Anaerobic Digestion of Waste Activated Sludge Digestion by the Addition of Zero Valent Iron. Water Res 2014, 52, 242–250. https://doi.org/10.1016/j.watres.2013.10.072.

(39) Golkowska, K.; Greger, M. Anaerobic Digestion of Maize and Cellulose under Thermophilic and Mesophilic Conditions – A Comparative Study. Biomass Bioenergy 2013, 56, 545–554. https://doi.org/https://doi.org/10.1016/j.biombioe.2013.05.029.

(40) Lee, Y. J.; Lee, D. J. Impact of Adding Metal Nanoparticles on Anaerobic Digestion Performance – A Review. Bioresour Technol 2019, 292 (July), 121926. https://doi.org/10.1016/j.biortech.2019.121926.

(41) Ripley, L. E.; Boyle, W. C.; Converse, J. C. Improved Alkalimetric Monitoring for Anaerobic Digestion of High-Strength Waste. 1986.

(42) Wiegant, W. M.; Hennink, M.; Lettinga, G. Separation of the Propionate Degradation to Improve the Efficiency of Thermophilic Anaerobic Treatment of Acidified Wastewaters. Water Res 1986, 20 (4), 517–524. https://doi.org/https://doi.org/10.1016/0043-1354(86)90202-2.

(43) Wang, Y.; Zhang, Y.; Wang, J.; Meng, L. Effects of Volatile Fatty Acid Concentrations on Methane Yield and Methanogenic Bacteria. Biomass Bioenergy 2009. https://doi.org/10.1016/j.biombioe.2009.01.007.

(44) Romero-Güiza, M. S.; Vila, J.; Mata-Alvarez, J.; Chimenos, J. M.; Astals, S. The Role of Additives on Anaerobic Digestion: A Review. Renewable and Sustainable Energy Reviews 2016, 58, 1486–1499. https://doi.org/10.1016/j.rser.2015.12.094.

(45) Zhang, W.; Wu, S.; Guo, J.; Zhou, J.; Dong, R. Performance and Kinetic Evaluation of Semi-Continuously Fed Anaerobic Digesters Treating Food Waste: Role of Trace Elements. Bioresour Technol 2015, 178, 297–305. https://doi.org/10.1016/j.biortech.2014.08.046.

(46) Owusu-Agyeman, I.; Plaza, E.; Cetecioglu, Z. Production of Volatile Fatty Acids through Co-Digestion of Sewage Sludge and External Organic Waste: Effect of Substrate Proportions and Long-Term Operation. Waste Management 2020, 112 (2020), 30–39. https://doi.org/10.1016/j.wasman.2020.05.027.

(47) Angenent, L. T.; Richter, H.; Buckel, W.; Spirito, C. M.; Steinbusch, K. J. J.; Plugge, C. M.; Strik, D. P. B. T. B.; Grootscholten, T. I. M.; Buisman, C. J. N.; Hamelers, H. V. M. Chain Elongation with Reactor Microbiomes: Open-Culture Biotechnology to Produce Biochemicals. Environ Sci Technol 2016, 50 (6), 2796–2810. https://doi.org/10.1021/acs.est.5b04847.

(48) de Souza Moraes, B.; Mary dos Santos, G.; Palladino Delforno, T.; Tadeu Fuess, L.; José da Silva, A. Enriched Microbial Consortia for Dark Fermentation of Sugarcane Vinasse towards Value-Added Short-Chain Organic Acids and Alcohol Production. J Biosci Bioeng 2019, 127 (5), 594–601. https://doi.org/10.1016/j.jbiosc.2018.10.008.

(49) Xie, S.; Li, X.; Wang, C.; Kulandaivelu, J.; Jiang, G. Enhanced Anaerobic Digestion of Primary Sludge with Additives: Performance and Mechanisms. Bioresour Technol 2020, 316 (July), 123970. https://doi.org/10.1016/j.biortech.2020.123970.

(50) Zhang, L.; Gong, X.; Wang, L.; Guo, K.; Cao, S.; Zhou, Y. Metagenomic Insights into the Effect of Thermal Hydrolysis Pre-Treatment on Microbial Community of an Anaerobic Digestion System. Science of the Total Environment 2021, 791, 148096. https://doi.org/10.1016/j.scitotenv.2021.148096.

(51) Wu, X.; Tian, Z.; Lv, Z.; Chen, Z.; Liu, Y.; Yong, X.; Zhou, J.; Xie, X.; Jia, H.; Wei, P. Effects of Copper Salts on Performance, Antibiotic Resistance Genes, and Microbial Community during Thermophilic Anaerobic Digestion of Swine Manure. Bioresour Technol 2020, 300 (30), 122728. https://doi.org/10.1016/j.biortech.2019.122728.

(52) Wang, P.; Wang, H.; Qiu, Y.; Ren, L.; Jiang, B. Microbial Characteristics in Anaerobic Digestion Process of Food Waste for Methane Production–A Review. Bioresour Technol 2018, 248, 29–36. https://doi.org/10.1016/j.biortech.2017.06.152.

(53) Guo, X.; Wang, C.; Sun, F.; Zhu, W.; Wu, W. A Comparison of Microbial Characteristics between the Thermophilic and Mesophilic Anaerobic Digesters Exposed to Elevated Food Waste Loadings. Bioresour Technol 2014, 152, 420–428. https://doi.org/10.1016/j.biortech.2013.11.012.

(54) Peng, X.; Börner, R. A.; Nges, I. A.; Liu, J. Impact of Bioaugmentation on Biochemical Methane Potential for Wheat Straw with Addition of Clostridium Cellulolyticum. Bioresour Technol 2014, 152, 567–571. https://doi.org/10.1016/j.biortech.2013.11.067.

(55) Kang, Y. R.; Su, Y.; Wang, J.; Chu, Y. X.; Tian, G.; He, R. Effects of Different Pretreatment Methods on Biogas Production and Microbial Community in Anaerobic Digestion of Wheat Straw. Environmental Science and Pollution Research 2021. https://doi.org/10.1007/s11356-021-14296-5.

(56) Schnürer, A.; Zellner, G.; Svensson, B. H. Mesophilic Syntrophic Acetate Oxidation during Methane Formation in Biogas Reactors. FEMS Microbiol Ecol 1999, 29 (3), 249– 261. https://doi.org/10.1016/S0168-6496(99)00016-1.

(57) Li, Z.; Wachemo, A. C.; Yuan, H.; Korai, R. M.; Li, X. Improving Methane Content and Yield from Rice Straw by Adding Extra Hydrogen into a Two-Stage Anaerobic Digestion System. Int J Hydrogen Energy 2020, 45 (6), 3739–3749. https://doi.org/10.1016/j.ijhydene.2019.07.235.

